# A pan-cancer signature of neutral tumor evolution

**DOI:** 10.1101/014894

**Authors:** Andrea Sottoriva, Trevor A. Graham

## Abstract

Despite extraordinary efforts to profile cancer genomes on a large scale, interpreting the vast amount of genomic data in the light of cancer evolution and in a clinically relevant manner remains challenging. Here we demonstrate that cancer next-generation sequencing data is dominated by the signature of growth governed by a power-law distribution of mutant allele frequencies. The power-law signature is common to multiple tumor types and is a consequence of the effectively-neutral evolutionary dynamics that underpin the evolution of a large proportion of cancers, giving rise to the abundance of mutations responsible for intra-tumor heterogeneity. Importantly, the law allows the measurement, in each individual cancer, of the *in vivo* mutation rate and the timing of mutations with remarkable precision. This result provides a new way to interpret cancer genomic data by considering the physics of tumor growth in a way that is both patient-specific and clinically relevant.

## Introduction

Understanding how a tumor evolves is clinically-valuable information: prognosis is determined by the future course of the evolutionary process^1,2^, and therapeutic response is controlled by the evolution of resistant subpopulations^3^. However, in humans the details of the evolutionary process have remain largely uncharacterized as longitudinal measurements are impractical, and studies are complicated by inter-patient variation^4^ and intra-tumor heterogeneity^5^. Moreover, the lack of a rigorous theoretical framework able to make predictions on existing data implies that results from cancer genomic profiling studies are often difficult to interpret. In particular, interpreting the allele frequency distribution reported by next-generation sequencing (NGS) data is problematic because of the lack of a formal model linking tumor evolution to the observed data.

Here we show that after the accumulation of tumor-driving alterations leading to the first malignant cell, the expansion of the cancer clone is dominated by effectively-neutral dynamics in a large proportion of cancers of different types and from different cohorts. These dynamics are described by a parameter-free analytical model of tumor expansion that predicts a power-law distribution of the allele frequencies of subclonal mutations in the tumor. We also demonstrate that reanalyzing cancer genome sequencing data in the light of neutral evolution reveals the mutation rate per division and the mutational timeline in each individual patient.

## Results

### Effectively-neutral cancer growth

Recently, we have validated the hypothesis that colorectal cancers (CRC) grow as a single expansion, populated by a large number of intermixed subclones^6^. These results predict that initially after malignant transformation, individual subclones grow at similar rates, despite showing distinct mutational patterns. In this context, it is the timing of occurrence of a new mutation that is the major determinant of its prevalence within the tumor, rather than clonal selection for that mutation.

Motivated by this latest evidence, here we developed a novel theoretical framework based on the concept of “effectively-neutral” evolutionary dynamics. The dynamics of neutral evolutionary processes have been studied extensively in the context of molecular evolution and population genetics^7-13^ and in mouse models of cancer^14^, but the presumption that subclone dynamics in human cancers are dominated by strong selection has meant that these ideas have not been applied to study cancer evolution. We derive a simple analytical result that specifically describes the distribution of mutant allele frequencies within a tumor that is measured by NGS data. Importantly, our model specifically considers the evolution of heterogeneous subclonal mutations.

A tumor is founded by a single cell that has already acquired a significant mutation burden^4^: these “pre-cancer” mutations will be borne by every cell in the growing tumor, and so become “public” or clonal. Within the growing malignancy, mutations that occur within different cell lineages represent “private” or subclonal mutations. During the growth of the cancer, on average new cells are born at rate *λ_b_* and die at rate *λ_d_*, corresponding to a net growth rate of *λ*=*λ_b_*–*λ_d_*. We note that the growth fraction may be the entire tumor (*λ_b_*=1) or only a small proportion of cells in active cell-cycle. The mean number of tumor cells as a function of time is then simply:

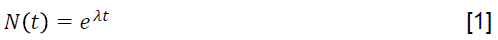

With a mutation rate *µ* per chromosome set per division and a ploidy *π* (number of chromosome sets in a cell), the average number of mutations occurring in a population of *N* cells is:

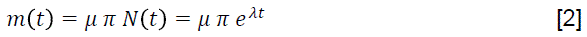

This equation is of limited use as neither *µ*, *λ* nor the cell cycle time can be measured directly in humans. Nevertheless, we do know that the allelic frequency of a newborn mutation at time *t* is the inverse of the number of chromosome sets in the population:

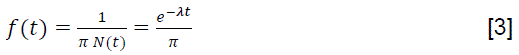

Importantly, in the absence of clonal selection (or indeed significant genetic drift), the allelic frequency (the relative fraction) of a mutation will remain constant during the expansion although the total number of cells with that mutation will increase. Solving for *t* and substituting into [2] gives an expression for the number of mutations *m(f)* at frequency *f*:

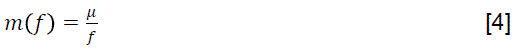

Hence, the distribution of mutant allele frequencies within a cancer is predicted to follow a 1/*f* power-law distribution with coefficient *µ*. The 1/*f* noise or *pink noise* is common in nature and found in several physical, biological and economic systems^15^. We note that this result is not only valid in the case of exponential expansion, but for any monotonic growth function *N(t)*. Critically, the distribution *m(f)* is naturally provided by next-generation sequencing data and importantly, it is independent of the parameters describing tumor growth, which are often immeasurable. To assess the accuracy of the model fit we use the cumulative distribution:

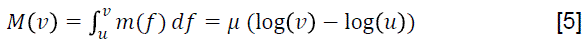

because it is less sensitive to “noise” in allelic frequency estimates inherent to NGS data. We note that in this model we do not consider public mutations as those have been accumulated in previous neoplastic stages for which the ancestral information has been lost. The model is therefore applied only to subclonal (private) mutations within a sample (the mutations describing malignant growth) whereas public mutations are filtered out. The range [u,v]=[0.25,0.1] for the integral in equation [5] is conservatively chosen to exclude public mutations, minimize the influence of copy number changes and account for the lower limit of detection in 50-100x depth sequencing, which is 5-10%^16^. We note that this detectability limit means that we are only able to study the initial growth of a tumor using “bulk” data (e.g. DNA from a large piece of tumor, rather than a single cell), since subclonal populations quickly fall below the detectability threshold as the tumor grows.

### Colorectal cancer evolution

The allelic frequency distribution of mutations in a tumor as measured by NGS whole-exome sequencing is shown in Figure 1A (data from ref ^6^). Mutations with high allelic frequency (>0.3) are likely to be public (clonal), however considerable information on the subclonal architecture of the tumor is encoded in lower frequencies mutations. The same data can be represented as the cumulative distribution in equation [5], where we focus only on the subclonal mutations captured by our model (Figure 1B). Remarkably, this follows precisely the log-linear distribution predicted by our model for this example in Figure 1B, as indicated by the high goodness of fit measure R^2^. In Figure 1C we considered our cohort of 6 multi-sampling CRCs^6^ and 106 TCGA colon adenocarcinomas^17^ that underwent whole-exome sequencing and contained at least 20 subclonal mutations (see Material and Methods for details). The latter were separated between tumors characterized by chromosomal instability (CIN) and microsatellite instability (MSI). The power-law distribution is remarkably well supported in both these independent cohorts, with 62/112 (55.3%) of the cases reporting a goodness of fit measure R^2^>0.95. These results confirm that in many CRCs, during the initial expansion of the tumor intra-tumor clonal dynamics are not dominated by strong selection but rather effectively-neutral evolution. We note that for a proportion of cancers with R^2^<0.95 this seems not to be the case. The poor fit in these samples may be due to lower sequencing quality, high normal contamination, or simply because these neoplasms do not develop according to an effectively-neutral evolution model.

**Figure 1.**
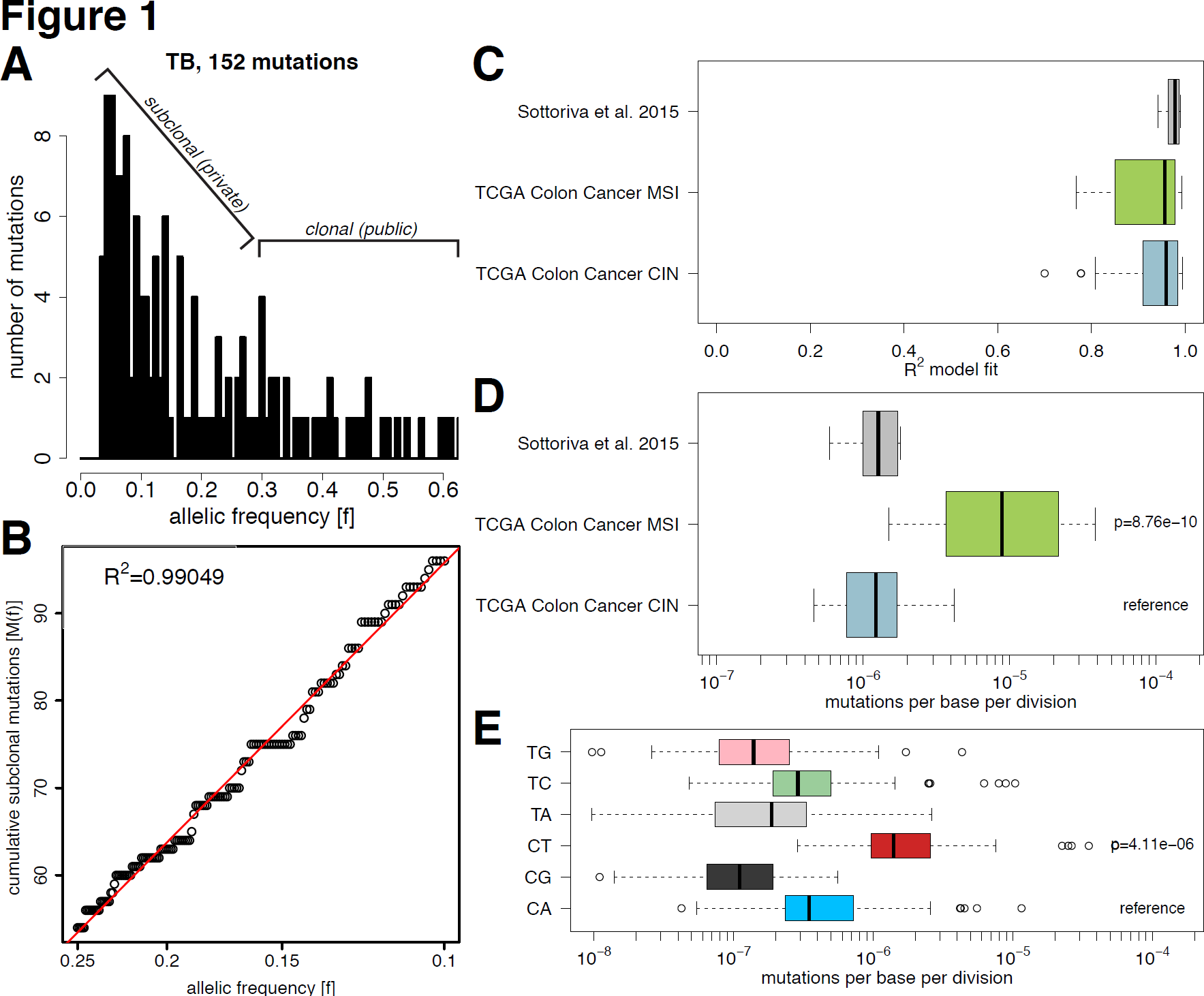
Effectively-neutral evolution is common in CRC and allows the measurement of the mutation rate per division in each tumor. **(A)** The output of NGS data such as whole-exome sequencing can be summarized as a histogram of the allele frequency of mutations, here represented for sample TB. Mutation with relatively high frequency (>0.3) are likely to be clonal (public), whereas low frequency mutations capture the tumor subclonal architecture. **(B)** The same data can be represented as the cumulative distribution M(f) of subclonal mutations. This was found to be log-linear with f, precisely as predicted by our effectively-neutral model. **(C)** R^2^ goodness of fit of our CRC cohort as well as a TCGA colon cancer cohort (n=112) grouped by CIN versus MSI confirmed that effectively-neutral evolution is common (55.3% with R^2^>0.95) **(D)** Measurements of the mutation rate showed that the CIN groups had median mutation rate of μ=1.22×10^−6^ whereas MSI tumors reported an almost 10-fold higher rate (median: μ=8.9×10^−6^, F-test: p=8.76×10^−10^). **(E)** Stratification by mutation type indicates that C>T mutations occur significantly at greater rate than other types.

Estimating the per-base mutation rate *µ* per division in human malignancies is challenging since direct measurements are not possible. Previous estimates critically depend on assumptions about the cell cycle time *T* and the mean tumor growth rate *λ*. Importantly, the analysis of tumor evolution using genomic data is dependent on quantification of the *total* mutational burden within the cancer^18-20^. Indeed, accurate measurement of all mutations within a cancer, including heterogeneous ones, is technically unfeasible since most mutations will be spatially isolated in small numbers of cells and so remain undetected by current sequencing assays^5^. However, in our model we are able to circumvent this issue by using only the subclonal mutations to measure the mutation rate per cell division in each individual patient, simply by fitting the gradient of the line given in equation [5]. We estimated the mutation rate in all samples with R^2^>0.95 and found an overall mutation rate of μ≈10^−7^−10^−6^ mutations per division in non-MSI cancers (Figure 1D). Expectedly, the mutation rate was almost 10-fold higher in the hypermutator MSI group (median: μ=8.9×10^−6^; F-test: p=8.76×10^−10^) with respect to the CIN group (median: μ=1.22×10^−6^) and our cohort (median: μ=1.14×10^−6^), which was comprised of all but one CIN tumors^6^. Different mutational types (e.g. transitions or transversions) are caused by particular mutational processes^21^, and so likely occur at different rates. Measuring the mutation rate of different mutation types revealed that C>T mutations occurred at median *µ_C>T_*=1.4×10^−6^, a rate nearly 10-fold higher than any other type of mutation (F-test: p=4.11×10^−6^; Figure 1E). We stratified according to CIN versus MSI and found that the mutation rate of each mutational type reflected the overall mutation rate for the group (Figure S1).

To further test effectively-neutral evolution, we contrasted the estimated mutation rate of synonymous mutations (unlikely to ever be under selection) versus the rate of missense and nonsense mutations (liable to experience selection). Although the latter group of mutations is more common than the former, after adjustment for the number of potentially synonymous or non-synonymous sites in the exome, the two rates were equivalent (t-test: p=0.38; Figure S2A), precisely as predicted by our model of neutral evolution. We also confirmed the robustness of our model to ongoing copy-number changes (Methods and Figure S2B,C).

### Pan-cancer analysis

We next applied our model to a large pan-cancer cohort of 883 exome-sequenced cancers from 14 tumor types. The fit was remarkably good across cancer types (Figure 2A) with 44.1% of the cases showing R^2^>0.95. Interestingly, we found that whereas several tumor types were clearly dominated by initial neutral evolution, such as colon, stomach, lung, prostate, cervical and bladder, others showed a consistently poorer fit, indicating that the clonal dynamics are typically not neutral in these malignancies. Other types displayed mixed dynamics, with some cases that were characterized by neutral evolution and some that were not. We can use the R^2^ value to identify cancer types that are more dominated by non-neutral dynamics like selective pressures, such as pancreatic, glioblastoma and renal. The latter is of particular interest as it displays convergent evolution likely driven by strong selective forces^22^, whereas convergent evolution was not found in lung cancer^23,24^, consistent with our results on the respective presence and absence of strong selection in the two cancer types.

**Figure 2.**
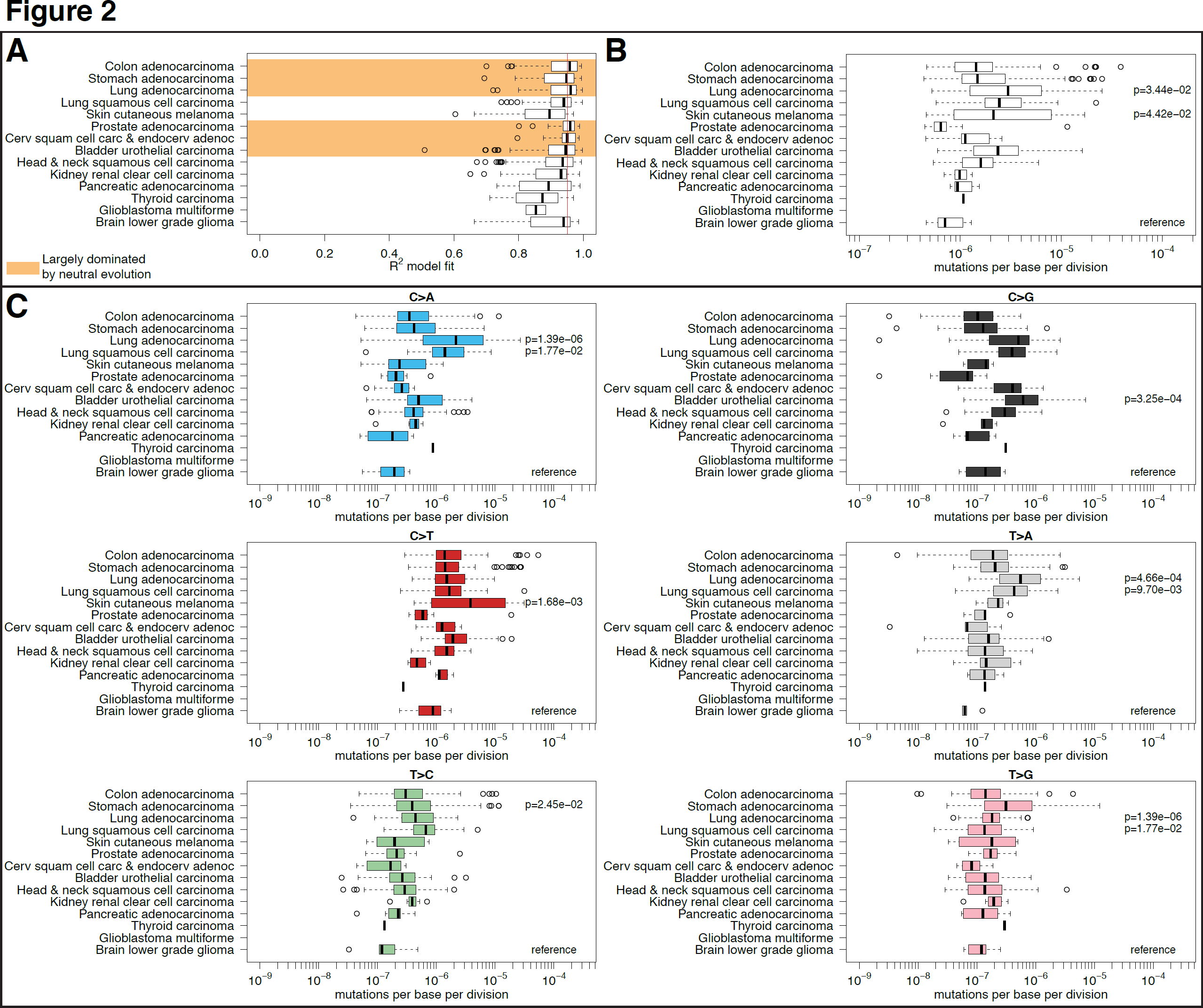
Pan-cancer effectively-neutral evolution and estimation of the mutation rate. **(A)** R^2^ values from 883 cancers of 14 different types supported our model in a large proportion of cases (44.1% of R^2^>0.95) and across the majority of cancer types, particularly colon, stomach, lung, prostate, cervical and bladder. On the contrary, pancreatic, glioblastoma and renal were characterized by non-neutral evolution. The other types displayed a mixed dynamics. **(B)** The highest mutation rate was found in lung cancer and melanoma, generally characterized by poor prognosis. Lower rates were found in thyroid, low grade glioma and prostate. **(C)** Stratification by mutation type revealed patterns consistent with exposure, such as higher C>A mutations in lung cancer due to tobacco and higher C>T mutations in melanoma from UV radiation.

The n=390 cancers with R^2^>0.95 were selected for mutation rate analysis, showing differences of more than an order of magnitude between types (Figure 2B). The highest mutation rates were observed in malignancies characterized by poor prognosis, such as lung adenocarcinoma (median: µ=2.98×10^−6^), lung squamous cell carcinoma (median: µ=2.44×10^−6^) and melanoma (median: µ=2.14×10^−6^). The lowest rates were registered for prostate (median µ=6.4×10^−7^) and low grade glioma (median µ=7.08×10^−7^). We stratified the mutation rates into different mutational types (Figure 2C) and found that C>A mutations occurred at a significantly higher rate in lung cancers, consistent with their causation by tobacco smoke^21^. C>T mutation rates were most consistent across cancer types, likely because of their association with normal replicative errors, as opposed to being caused by a particular stochastically-arising defect in DNA replication or repair^25^. Melanoma had the highest rate of C>T mutations, as expected from the skin’s exposure to UV radiation.

Importantly, equation [2] demonstrates that there is a relationship between the mutation rate and the growth rate of the tumor, meaning that our estimate of the mutation rate is dependent on the unknown symmetric cell division rate that drives cancer growth. Consequently, if symmetric divisions are rare, we acknowledge that this may lead to an overestimation of the real mutation rate per division.

### Mutational timelines

Recognition that the initial growth of a tumor follows effectively neutral dynamics allows estimation of the size of the tumor when a mutation with frequency f occurred:

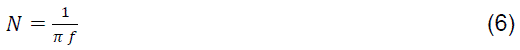

Figure 3A,B shows the decomposition of the mutational timeline for two illustrative cases, namely sample TB from^6^ and sample TCGA-AA-3712 from^17^. Previous estimates of this mutational timeline relied on cross-sectional data^26-29^ which are compromised by the extensive heterogeneity, whereas multi-region profiling approaches are instead accurate but expensive and laborious^22,30,31^. Using our formal model of cancer evolution this timeline information becomes accessible from routinely available genomic data. We note that only the order of initial mutations can be deciphered by our method, because of the difficulty in detecting rare mutations discussed above (Figure 4A). We found that classical CRC driver alterations, such as in the *APC*, *KRAS* and *TP53* genes were indeed present in the first malignant cell (likely because they accumulated during previous neoplastic stages). This confirms what we previously reported using single-gland mutational profiling where all these drivers, when present, were found in all glands^6^. However, we also found that when we considered a more extended list of putative drivers, many occurred after the initial seeding of the malignancy implying they had not experienced strong selection.

**Figure 3.**
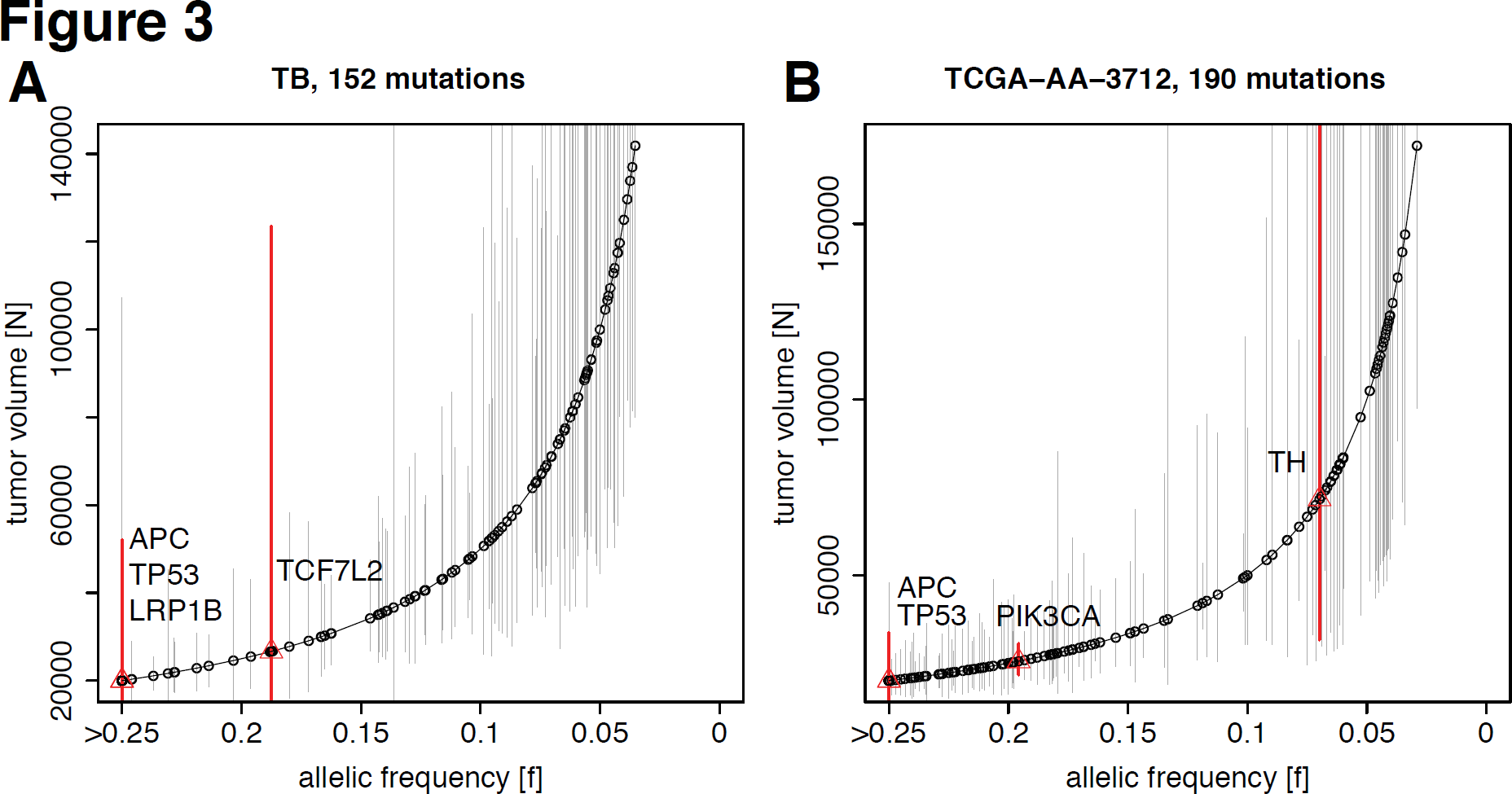
Reconstruction of the mutational timeline in each patient. The frequency of a mutation within the tumor predicts the size of the tumor when the mutation occurred. **(A,B)** The deconvolution of the mutational timeline is illustrated for sample TB and TCGA-AA-3712, in which whereas established CRC drivers (APC, KRAS, TP53) were found to be present from the first malignant cell, several non validated putative drivers were mutated after malignant seeding, despite the underlying neutral dynamics. This suggests that some of these candidate alterations may not be fundamental drivers of growth in all samples. Confidence intervals are calculated using a binomial test on the number of variant reads versus the depth of coverage for each mutation.

**Figure 4.**
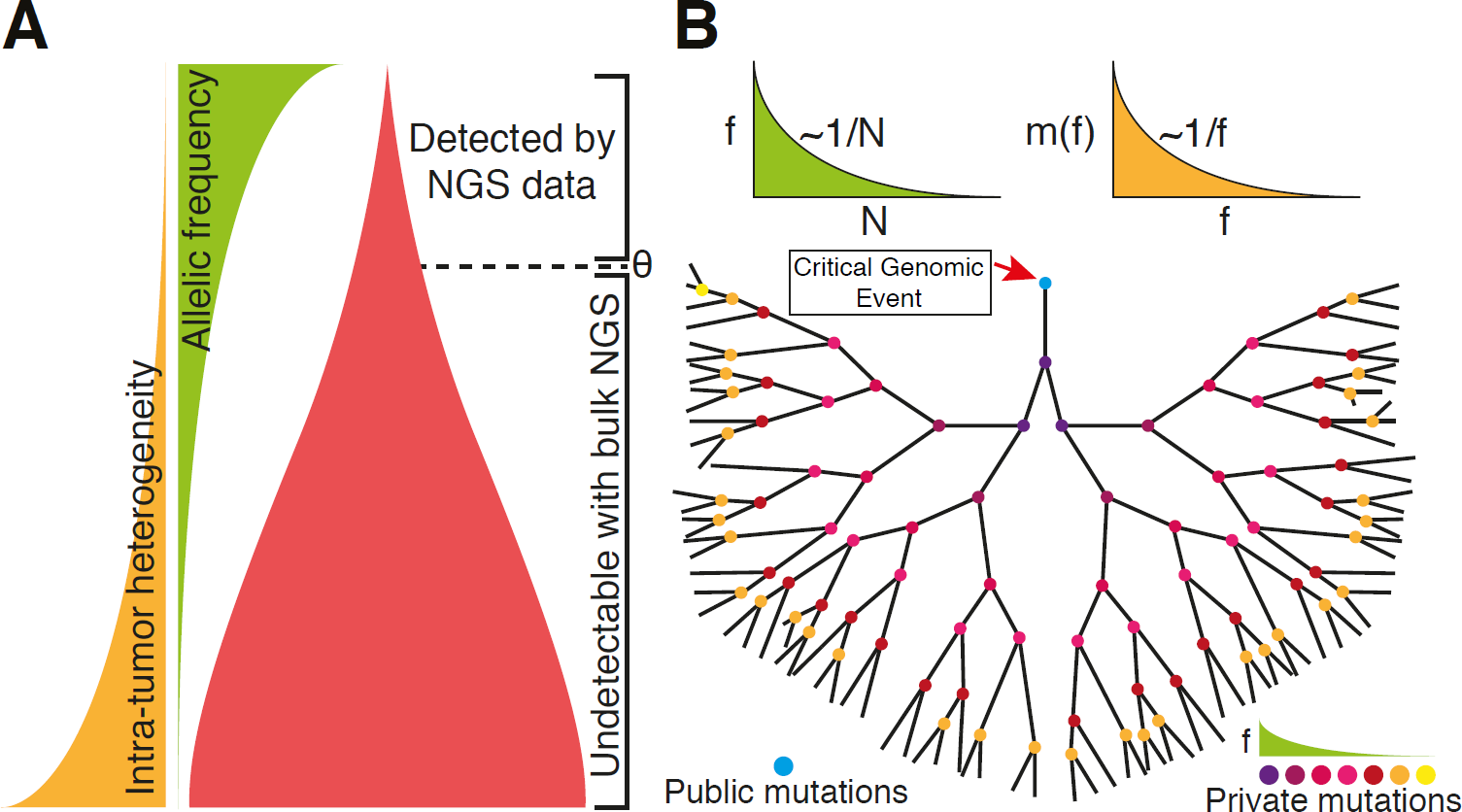
The neutral evolutionary model. **(A)** Current sequencing technologies using bulk samples detect only mutations occurring during the initial tumor growth because the allelic frequency of new mutations quickly falls below the detection limit θ as the tumor grows. Thus, low-frequency late-arising mutations go undetected using bulk sample profiling. **(B)** After the accumulation of genomic alterations, the cancer expansion is likely triggered by a single critical genomic event (the accumulation of a “full house” of genomic changes) followed by neutral evolution that generates a large number of new mutations in ever smaller subclones. In this context, the allelic frequency of mutations f(t) follows a 1/N distribution and, consequently, the accumulation of mutations m(f) follows a characteristic 1/f distribution. Deviations from this law indicate instead the presence of selection or significant drift.

## Discussion

Understanding the evolutionary dynamics of subclones within human cancers is challenging because longitudinal observations are unfeasible. Here we have demonstrated that the initial growth of cancer is often dominated by neutral evolutionary dynamics, an observation that is consistent across 14 cancer types. For the first time we present a mathematical law of mutational accumulation during cancer growth that predicts mutational patterns routinely reported using NGS. This result suggests that following the critical genomic event(s) that initiate clonal expansion the subsequent sub-clone evolution is effectively neutral and so the frequency (f) of private mutations is characterized by a 1/f distribution (Figure 4B). In this context, it is possible that the microenvironment indeed does not play a fundamental role in driving the initial subclone evolution, since despite genetic differences between cells and the presumable differences in the microenvironment they reside in, the overall pattern of evolution is effectively neutral. Further, it is tempting to suggest that eventual adaptation to distinct microenvironments may be due to cancer cell plasticity, rather than clonal selection: the lack of clonal selection observed suggests that the original genomic events that triggered malignant growth may inherently cause a plastic phenotype and so facilitate adaption to different microenvironments without requiring clonal evolution. We note that some cancer types were more dominated by neutral evolution than others. We speculate that such evolutionary dynamics may be driven by the cellular architecture of the tumor (e.g. glandular structures that limit the effects of selection) and/or the anatomical location of the malignancy (e.g. growing in a lumen versus growing in a highly confined space). Despite the evidence for lack of natural selection during malignant growth, eventual treatment is likely to change the rules of the game and strongly select for treatment resistant clones. The same may happen in the context of the purported evolutionary bottleneck preceding metastatic dissemination.

Importantly, the realization that the within-tumor clonal dynamics are effectively-neutral means that the *in vivo* mutation rate per division can be directly measured in a sample. This parameter plays a key role in cancer evolution, progression and treatment resistance. Importantly, these measurements can be performed in a patient-specific manner and so may be useful for prognostication and the personalization of therapy. Recognizing that the initial growth of a neoplasm is dominated by effectively-neutral clonal dynamics provides an analytically tractable and rigorous method to study cancer evolution and gain clinically relevant insight from commonly available genomic data.

## Online Methods

The processing of exome-sequencing data from^6^ and TCGA^17^ involved PCR duplicate removal and variant calling on matched-normal pairs using Mutect^16^. A mutation was considered if the depth of coverage was ≥ 10 and at least 3 reads supported the variant. Mutations that aligned to a more than one genomic location were discarded. Non-CRCs in the TCGA had mutations called using Mutect according to the pipeline described in^32^. Microsatellite instability in the TCGA colon cancer samples was called using MSIsensor^33^.

To fit our model to allele frequency data we considered only variants with an allele frequency in the range [u,v]. The low boundary v is due to the fact that we have a limit in NGS data for the reliable detectability of low-frequency mutations in the order of 5-10%^16^. The high boundary u is necessary to consider only subclonal alterations and not public mutations that were present in the first transformed cell. In the case of diploid tumors, this threshold is 0.5 (mutations with 50% allelic frequency are heterozygous public or *clonal*), in the case of triploid tumors, this threshold drops to 0.33. For all samples we used a boundary of [0.1-0.25] to account only for reliably called mutations and normal contamination in the samples. All the samples considered in this study had a minimum of 20 reliably called private mutations within the fit boundary. Once these conditions were met in a sample, equation [5] was used to perform the fit as illustrated in Figure 1B.

Ongoing copy-number change (allelic deletion or duplication) can alter the frequency of a variant in a manner that is not described by equation [4]. We assessed the impact of copy-number alterations (CNAs) on our estimates of the mutation rate within the TCGA colorectal cancer samples by using the paired publically available segmented SNP-array data to exclude somatic mutations that fell within regions of CNA. CNVs were identified having an absolute log-R-ratio>0.5, and the model fit was performed only on diploid regions of the genome. Figure S2B shows that the fit of the model is not perturbed by copy number alterations since, despite the reduced number of mutations per sample, the goodness of fit is still remarkably high. Furthermore, the estimates of the mutation rate based only on the remaining variants (e.g. variants located in diploid regions of the genome) were in excellent agreement were estimates based upon all variants (Figure S2C).

## Contributions

AS and TG jointly contributed to the development of the model, the analysis of the data and the interpretation of the results.

## Acknowledgements

AS is supported by The Chris Rokos Fellowship in Evolution and Cancer. This work was supported by the Wellcome Trust [105104/Z/14/Z]. We thank Prof. Darryl Shibata for the fruitful discussion. We would like to that Noemi Andor (Stanford University) for supplying mutation calls for the TCGA data.

---

**Figure S1.**
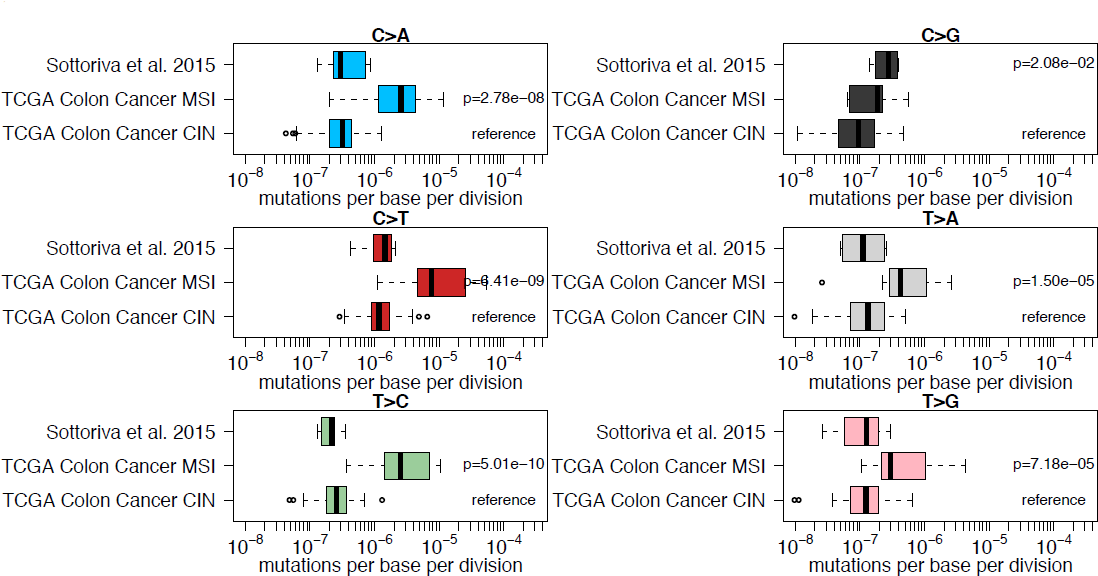
Rates of different mutation types in CRC. As for the overall mutation rate (Figure 2B), all mutation types apart from C>G were significantly higher in the MSI group.

**Figure S2.**
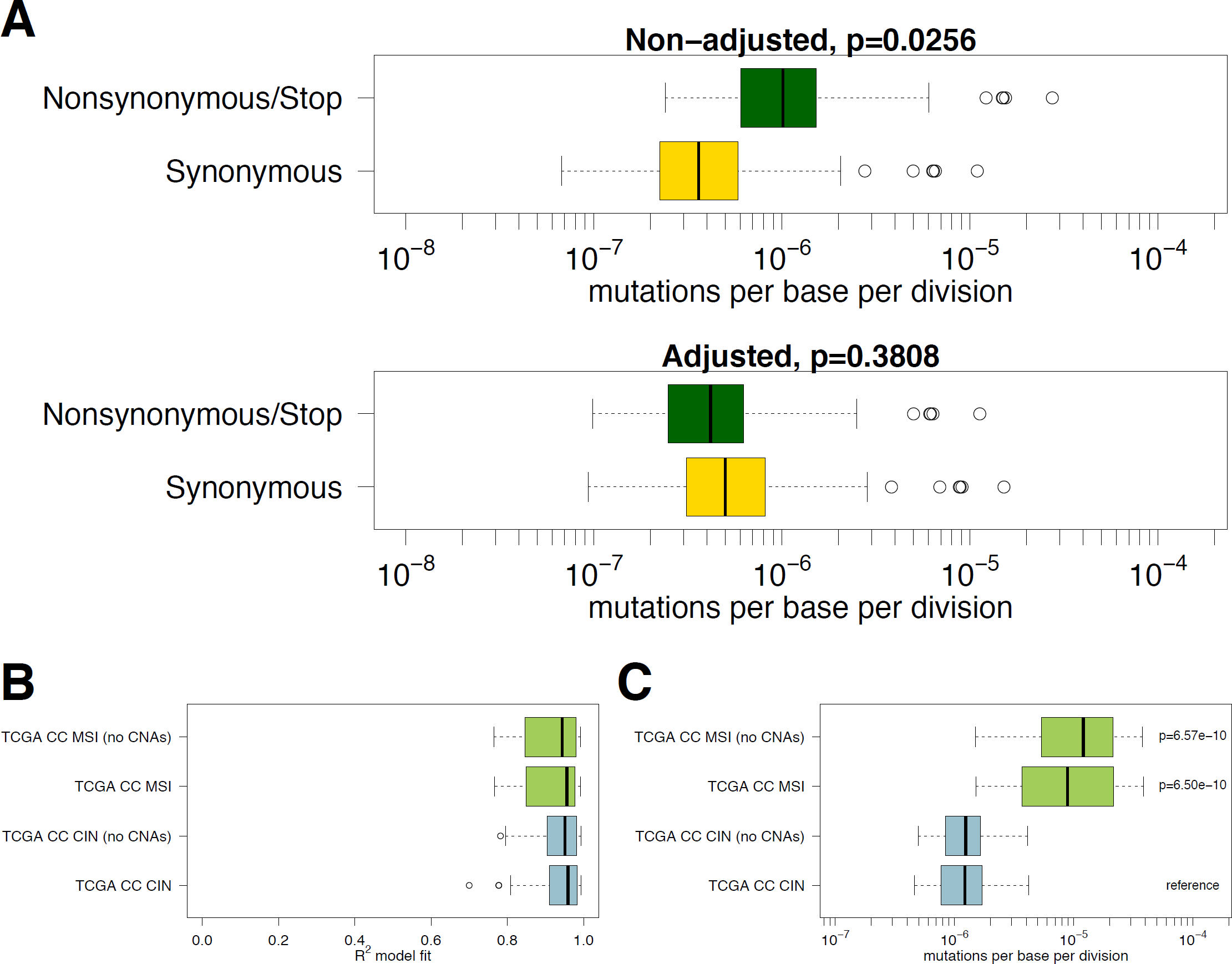
Neutrality is further validated using synonymous versus nonsynonymous/stop mutations and is robust to copy number changes. (A) A random base change within a codon is more likely to result in a nonsynonymous or stopgain mutation than a synonymous mutation; hence we expect the mutation rate per division of non-synonymous mutations to be higher than for synonymous mutations. However, the neutral model predicts that when adjusting for the total number of possible non-synonymous versus synonymous sites, these two rates should be the same. This is indeed true (t-test; p=0.38), as shown for the colon cancer samples reported in Figure 1, thus providing further validation for our neutral model of cancer growth. **(B)** Using SNP arrays paired to the exome sequenced samples we subtracted from the analysis those mutations that fell within regions of the genome with altered copy number. The consistent high values of goodness of fit demonstrate that our model is robust to confounding copy number changes. **(C)** Estimating the mutation rates using only the mutations in copy number devoid regions yields the same results, confirming the robustness of our approach.

